# An integrase clade that repeatedly targets prophage late genes, yielding helper-embedded satellites

**DOI:** 10.1101/2022.07.18.500453

**Authors:** Dario Tommasini, Catherine M. Mageeney, Kelly P. Williams

**Affiliations:** Sandia National Laboratories, Systems Biology, 7011 East Avenue Livermore, CA

**Keywords:** prophage, PICI, bioinformatics, genomic island

## Abstract

Satellites are mobile genetic elements that rely on helper phages for their mobilization. The many known satellite-helper interactions are trans-regulatory, with gene products from one partner modulating the nucleic acid or protein activities of the other. We discovered a satellite type with a more intimate cis-regulatory configuration: integrated within, and co-oriented with, a late gene of its lambdoid helper prophage. This helper-embedded satellite (HES) configuration would delay expression of the interrupted helper late gene until the satellite excises; it also offers potential passive components to both HES replication and late transcription, driven by the helper. Induction of a helper-satellite composite was monitored; precise excision of the entire composite was observed, followed by its replication, and the excision of the satellite from it. We mapped 491 HESs to one of 14 sites in cognates of phage lambda late genes A, B, C, E, V, T, H, L and J. The associated integrases form a single phylogenetic clade with subclades respecting the 14 site groups, while the *attP* attachment site regions contained a new doubled DNA sequence motif. This clade thus exhibits a repeated tropism for prophage late genes as it develops new integration sites. HESs bear close genomic similarities to gram-negative phage-induced chromosomal islands (PICIs, of which we found many more integrated into *fis* and *hpt* genes). We describe four ordered zones in a general HES/PICI genome organization: an integration zone encoding integrase and AlpA, a Bro zone encoding members of the Bro-N network of domain-swapping DNA-interactive proteins and immunity repressor RNAs, a replication zone, and a late zone in which clusters as large as 18 consecutive helper late genes have been captured. Like the late zone, the Bro zone is dynamic, perhaps due to activity of the Bro proteins themselves.

## INTRODUCTION

So, Nat’ralists observe, a Flea
Hath smaller Fleas that on him prey,
And these have smaller yet to bite ‘em,
And so proceed *ad infinitum*
–*Swift, On Poetry: A Rapsody*

Bacteriophage satellites are small mobile genetic elements that require specific helper phages for their mobilization^1–5^. A large group of satellites with shared gene content includes the classical satellite P4, the gram-negative and gram-positive lineages of the phage-induced chromosomal islands (PICIs), and PICI-like elements (PLEs) found in *Vibrio cholerae* ^6–14^. Another *Vibrio* satellite, TLCϕ, differs from the above by its use of a filamentous, rather than tailed, helper phage ^15^. Satellites exploit virion structure and assembly functions of the helper, at the expense of helper virion yields. They can adapt to their helpers, exhibiting genetic interactions that can be extensive and reciprocal. For example, P4 can derepress early genes and activate late genes of its helper P2, and respond in the same ways to P2-encoded regulators^1^.

Although at least some satellites can optionally replicate in circular form like plasmids, they are known for integrating into the bacterial chromosome, and can thus be classified as genomic islands (GIs), as are their helper prophages. Each GI usually encodes an integrase that specifies its chromosomal integration (attachment) site, *attB*, and the corresponding attachment site in the GI, *attP*. Integration converts *attB* and *attP* into the GI-flanking recombinant site pair *attL* and *attR*. The set of four *att* sites typically share a block of sequence identity of at least 7 bp, within which DNA strand crossover occurs during recombination. Our software TIGER^16^ detects and maps GIs in bacterial chromosomes, with sufficient precision to delineate the *attL* and *attR* sites. TIGER revealed a nested pattern in one *E. coli* genome, where a small element, 11capE, was integrated within a larger prophage, 50icd. A satellite/helper relationship was suggested by close genomic similarities between the 11capE/50icd pair and satellite/helper pairs among the gram-negative PICIs (GN-PICIs).

Despite their recent recognition as a family, some interesting mechanistic aspects of the GN-PICIs have emerged^6, 7^. Three conserved genes are essential for induction and PICI transfer, *int* (integrase), *alpA* (a DNA-binding regulator), and *pri* (the replication primase); these, and most of their other genes, are unidirectional. AlpA activates its own promoter, and may^17^ or may not^6^ activate the upstream *int* promoter; it is also known as a recombination directionality factor (excisionase)^18^. An origin of replication (*ori*) is found downstream of *pri*. A newly discovered PICI-encoded protein, RppA, directs helper terminase to PICI DNA for packaging into virions, at the expense of helper DNA packaging. GN-PICIs can carry phage late genes, most often a gene for the major capsid protein, which is present in 21 of 29 reference PICIs; however, such PICI capsid genes are not required for helper-assisted PICI transfer^6^. GN-PICI genomes are small (10-20 kbp), yet they are packaged into longer, phage-sized (~45 kbp) DNA concatemers in the virion, unlike some other satellites; P4 and some gram-positive PICIs are packaged in single-genome DNA lengths in smaller virion heads.

Here, we monitored induction of the 11capE/50icd helper/satellite, observing excision of the composite, followed by replication of the composite and excision of the satellite. Bioinformatic analyses revealed numerous relatives of the 11capE satellite among the Enterobacterales, each integrated within one of nine prophage late genes, and co-oriented with the late gene. This configuration, which we term helper-embedded satellite (HES), implies a new dimension to satellite/helper relationships.

## MATERIALS AND METHODS

### Bacterial strain

The 11capE HES-bearing strain, *Escherichia coli* NRG857c (abbreviated here Eco567), was kindly supplied by Alfredo Torres (U. Texas Medical Branch).

### Bioinformatic identification of HESs, PICIs and prophages

Our founding HES, 11capE, its surrounding unit, 61icd, and other prophages were identified and mapped in the chromosome (CP001855.1) of Eco567 using TIGER^16^ software as previously described.

To examine relatives of 11capE, its integrase sequence was used as a TBLASTN query against 165,778 proteobacterial genomes of reasonable quality (N50 > 10000; < 300 contigs) downloaded from GenBank in July 2019, that were split into 7 databases as follows: 58711 *Salmonella*, 27909 other Enterobacteriaceae, 2499 other Enterobacterales, 24897 other Gammaproteobacteria, 30634 *Campylobacter*, 1945 other Epsilonproteobacteria, and 19183 other Proteobacteria. The top five hits in each database were examined for phage gene surroundings, which were found only among the Enterobacterales. All the non-Enterobacterales integrase matches scored below 400 bits; this cutoff was applied, leaving 5068 enterobacterial integrase genes (*int*s) for further examination. Regions of ~22 kbp (the *int*-matched segment with flanks of 1 kpb upstream and 20 kbp downstream) were taken, rejecting *int* regions containing an ambiguous base or truncated due to reaching a contig end, leaving 3490 *int* regions. During TIGER processing it became clear that non-HES PICIs integrated into the *fis* gene were an abundant artifact; jobs were cancelled for 230 yet-unprocessed regions containing a *fis* gene. TIGER yielded 3177 satellite calls among the 3260 treated regions, although 14 of these required curation (splitting) of the annotation of an integration-generated gene fusion between *int* and the upstream prophage late gene fragment. For the 83 initial failures, application of TIGER to the entire genome mapped a satellite in 18 cases elongated beyond the ~22-kbp region and in 14 cases where normal-sized satellites had been missed for the isolated region. For eight additional cases, a HES was mapped manually by inspection of split late genes in the region; TIGER failure in these cases was explained by an insertion sequence very close to *attR* (one case) and by occurrence in the genus *Arsenophonus* which only had 7 reference genomes in the database (five cases). This left 43 regions where at least one terminus of the satellite could not be mapped; for 34 of these, the target gene could be guessed based on the coding sequence upstream of the integrase gene, but *attR* could not be identified, while nine remained fully unmapped, often associated with transposase genes near *int* that suggested chromosomal rearrangements. Three calls were recognized as two-PICI tandems that we split into their components (Supp. Figure 1).

For all genomes with a fully mapped HES, TIGER and Islander were run for the entire genome, delineating 208 HES-surrounding lambdoid helper prophages, and 24 *E. coli* HES-surrounding PICIs with the HES integrated into the PICI late gene site C:CLP_protease: 110. For all these, orientation was assigned as that of the HES, and the hypothetical form with the HES excised was taken for further analysis. This yielded a total of 3452 newly mapped GIs, comprising 208 prophages, 2725 PICIs, and 519 HESs (Supp Table 1). We added reference GIs to our study set – 28 reference GN-PICIs^6^, their one known helper (Eco160.55.icd), P4 and its helper P2, and lambda, for a total of 3484 study GIs. Reanalysis of reference PICIs by TIGER, Islander and inspection remapped 15 or more to tRNA genes (Supp Table 1). To summarize taxonomy, all GIs studied here are from 33 genera of the Enterobacteriaceae, except for reference PICIs from the Pasteurellaceae genera *Aggregatibacter, Necropsobacter and Pasteurella*.

### Gene content

The 3484 study GIs were annotated using the Tater pipeline^19^, which supplements the Prokka annotations of Prodigal CDS calls with Pfam HMM hits, and carefully annotates integrase genes.

Initial clustering of proteins revealed that Bro network proteins were improperly aggregated, so they were treated specially. Primary hits to 10 Pfam HMMs of the Bro network were found among the GIs: ANT, AntA, Bro-N, KilA-N, ORF11CD3, ORF6N, P22_AR_C, P22_AR_N, Phage_ASH, and Phage_pRha. To account for possible new Bro domains, the flanking peptide sequences of these hits were used as BLASTP queries (evalue 0.00001) to find additional candidate Bro proteins. These proteins sequences were split into their hit and unhit segments; the segments were clustered using MCL^20^ with an inflation parameter of 1.1), and aligned using MAFFT in “auto” mode. The alignments were curated manually, including both breaking clusters into MAFFT -aligned subsets and splitting segments further based on closely related blocks (Supp File 1). Each of the 144 groups of candidate Bro segments was designated with a “b” followed by a serial number. To prevent spread into non-Bro proteins through occasional frameshifts, we rejected 24 rare Bro segments found more frequently outside than inside the original candidate Bro proteins. Since Prodigal occasionally may have miscalled ORFs, each segment was used as a TBLASTN query (cutoff 30 bits) against the GI genomes, identifying candidate Bro CDS, each labeled according to its sequence of segments.

Annotations in raw tater GFF outputs that shared a stop codon with a Bro protein were removed and Bro protein annotations were merged in. TBLASTN with AlpA queries showed that Prodigal had failed to call the AlpA reading frame in 5 satellites, each corrected. With the Bro proteins and the integrase family (Int) treated, the remaining 10334 unique proteins were treated by all vs. all BLASTP, with a cutoff of 30 bits, and clustered using MCL^20^ with an inflation parameter of 1.4, producing 1395 families (682 singletons). A large family containing hits to Pfams Pox_D5, D5_N, DUF927, Prim-Pol, or Prim_Zn_Ribbon was renamed “Primase”. Otherwise each family was named according to its most frequent Pfam hit, or if none, was considered hypothetical and named ‘H’ with a serial number (847 families). The family names of all encoded proteins were concatenated in genomic order, preceded with a ‘-’ symbol if oriented backwards, to produce 646 unique gene profiles, 168 for prophages and 480 for satellites.

The 26 families with either “tail” or “holin” in their names (all with late gene functions) were used as seeds to collect strings of continuous co-oriented genes from the prophage profiles. Such strings do not always perfectly separate early from late genes. For example, late genes lie downstream of and co-oriented with the lambda early gene Q, and an operon of two non-seed P2 late genes is transcribed oppositely to the other late genes. However this simple approach was mostly effective at separating known early and late gene families for both lambda and P2 (Supp. Figure 2) except for low-abundance families in the situations mentioned above. An additional 32 families enriched in late strings and associated with late phage functions were added to the seed and known late gene families, for a set of 93 high-confidence families categorized as Late, that account for 39.3 % of all prophage profile genes. Other categories applied were Early (the known early gene families of lambda and P2 only), Integration (Int, Phage_AlpA and HTH_17 families), Replication (Primase, SSB and DUF5906 families), Bro (the Bro proteins), Hypothetical and Other.

Genome annotations and BLASTN comparisons were visualized according to gene content using EasyFig^21^.

### Phylogenetic analysis

The 3494 proteins hit by the Phage_integrase Pfam HMM, from the 3484 GIs treated for gene content as above (where 9 HESs or PICIs contained secondary or tertiary integrase genes outside the usual integration module), were combined with those of the 43 guessed or unmapped satellites and the 24 integrase sequence fragments used to seed the Pfam Phage_integrase. Sequences were aligned using ‘mafft --auto’ and built into a phylogenetic tree using ‘FastTreeDbl -lg’.

### Prophage induction

Three replicates were performed simultaneously, each started from a different colony of Eco567 grown as an overnight culture, which was diluted 100-fold and grown in LB to mid log phase (OD_600_ = 0.4-0.5). Mitomycin C was added to 1 μg/ml and 1-ml samples were taken at 0, 0.5, 1, 2, and 3 hr after treatment. Samples were centrifuged at 5000 x g for 2 min. The supernatant was removed and passed through a 0.22 μm filter. Genomic DNA was isolated from the pellet with the DNeasy Blood & Tissue Kit (Qiagen). Genomic DNA yields were quantitated using Qubit Flex Fluorometer (Thermofisher).

### Quantitative real-time PCR (qPCR)

PCR products (Supp. Figure 3) were prepared using QIAquick PCR Purification Kit (Qiagen) and quantitated using Qubit Flex Fluorometer (Thermofisher), as standards for each of the eight *att* site forms of interest (*attB, attP, attL*, and *attR* for both 61icd and 11capE), using the primers listed in Supp.Table 2. Genomic DNA samples of 40-60 ng were prepared, along with a hundred-fold dilution series (100 pg, 1.0 pg, 10 fg, and 100 ag) of standard curve samples. PCR reactions were performed in PowerUp SYBR Green Master Mix (Applied Biosystems) with each primer at 500 nM using a three-step amplification protocol (58°C annealing) in a CFX96 Touch Real-Time PCR Detection System (Bio-Rad).

The slope and intercept of each molar standard curve was taken, and the following formula used to convert from Ct value to copy number in the genomic DNA sample: copies = 10^(Ct-intercept)/slope^. All reaction efficiencies were above 88% and all r^2^ values for standard curves were above 0.98. The total copy number of *icd* regions was taken as the *icd attB* counts summed with the average of *icd attL* and *attR* counts. Finally, *attB* and *attP* counts were reported after normalizing to *icd* counts.

### Quantitation of *att* sites through deep sequencing

Deep sequencing was employed as a second method of measuring *att* sites, through counting of reads from each *att* site type^22^. Sequencing libraries were prepared from the genomic DNA samples of one of the induction experiment replicates using the Nextera DNA Flex library prep kit with the Nextera DNA CD Index kit (Illumina). Libraries were quantitated using the Qubit high-sensitivity DNA assay kit (Thermofisher Scientific) and pooled in equal quantity to form a multiplexed final library, requantitating with Qubit and confirming quality with the Agilent bioanalyzer using high sensitivity DNA chip (Agilent Biotechnology). The final library was sequenced using Illumina technology on a NextSeq 500 sequencer with a high-output 150-bp single-end read kit.

Sequencing reads were quality filtered using BBDuk (http://jgi.doe.gov/data-and-tools/bb-tools/). The quality filtered reads were analyzed with Juxtaposer^22^ to identify mobile genetic elements and AttCt^22^ software to determine *attL, attR, attB*, and *attP* read counts for each GI. Probes used by AttCt for each predicted genomic island are listed in Supp Table 3. The *attB* and *attP* read counts were normalized to *icd* region (or *torS* region) counts as described above for qPCR.

## RESULTS

### A nested pair of genomic islands

When considering a possible island in a genome, if a reference genome can be found that represents a precise deletion of the island, this constitutes support for the island, and maps it. Stronger support comes from finding more such reference genomes. This comparative approach is automated by our recently developed computer program TIGER^16^, notable for its high precision of mapping, sufficient to delineate the attachment sites (*att*) of integration. In a search for GIs that encode a homolog of the same chromosomal gene into which they integrate^16^, we found in *Escherichia coli* NRG857c a small GI, Eco567.11capE (Supp. Table 1). Our GI nomenclature combines an abbreviation and serial number for the strain with the size and the gene target of the island. 11capE is integrated within (and disrupting) a phage capsid gene, yet bears its own capsid gene. 11capE was nested within another GI call, 61icd, that TIGER identified as a prophage. The 61icd integration site in the *icd* gene is a well-known prophage integration site in *E. coli*^23^. We hypothesized that 61icd is a two-part composite, with the 11capE component integrated into the capsid gene of a putative 50-kbp prophage that we term 50icd. Sequence comparison of the left and right attachment sites (*attL* and *attR*) of 11capE (Figure 1) reveals a 10-bp sequence identity block where integration occurred. The 50icd capsid gene is a very close relative of the major capsid gene E of phage lambda, and the capsid gene within 11capE is from the same family, but extensive sequence differences in the region corresponding to the attachment site would likely prevent 11capE self-integration.

**Figure 1.**
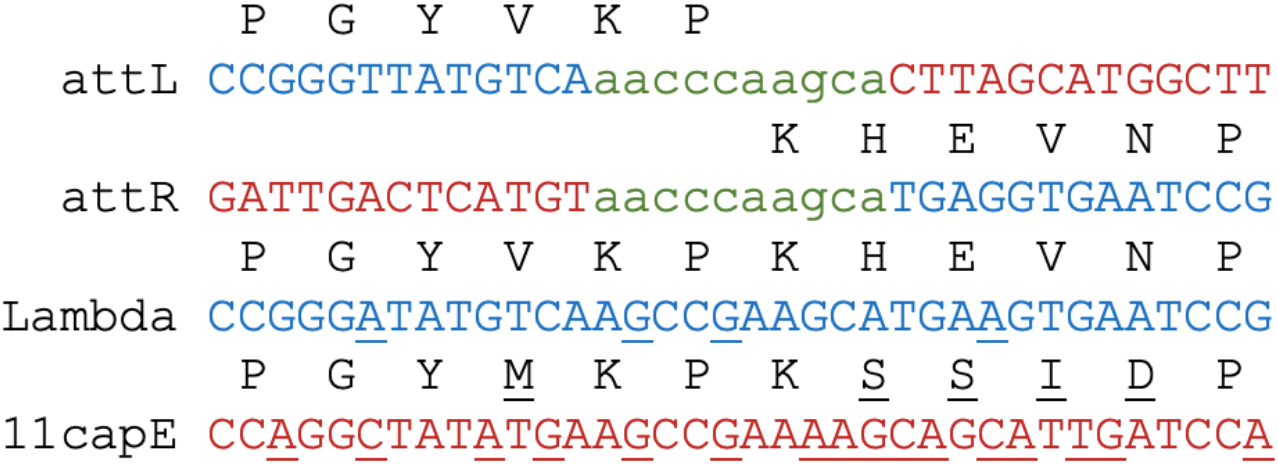
Integration site of 11capE in the capsid gene of prophage 50icd. The left and right *att* site sequences of 11capE in the composite 61icd are shown with encoded capsid sequence above; 50icd sequence in blue, 11capE sequence in red, and the *attL/attR* sequence identity block in green lower case. Corresponding segments are shown from the major capsid gene E of phage lambda and the homolog encoded within 11capE, with sequence differences from *attL/attR* underlined.

11capE shares the size and gene organization of the recently discovered gram-negative PICIs, and some sequence similarity, as shown in Fig. 2 for the emerging model PICI^6, 7^, EcCICFT073. The helper prophages of 11capE and EcCICFT073 likewise show extensive similarity to each other (in this case sharing the same chromosomal integration site in the *icd* gene), and to phage lambda (Figure 2B). Whereas other PICIs are integrated into chromosomal genes apart from their helpers, allowing interaction only *in trans*, 11capE is not only “phage-induced”, it is phage-embedded. Integration within its helper prophage allows a unique cis-regulatory relationship between satellite and helper that we discuss below. Clearly, expression of the 50icd capsid gene cannot occur until 11capE excises. Furthermore, both late transcription and replication of the helper should also propagate to the integrated satellite. We term this new nested configuration a helper-embedded satellite (HES).

**Figure 2.**
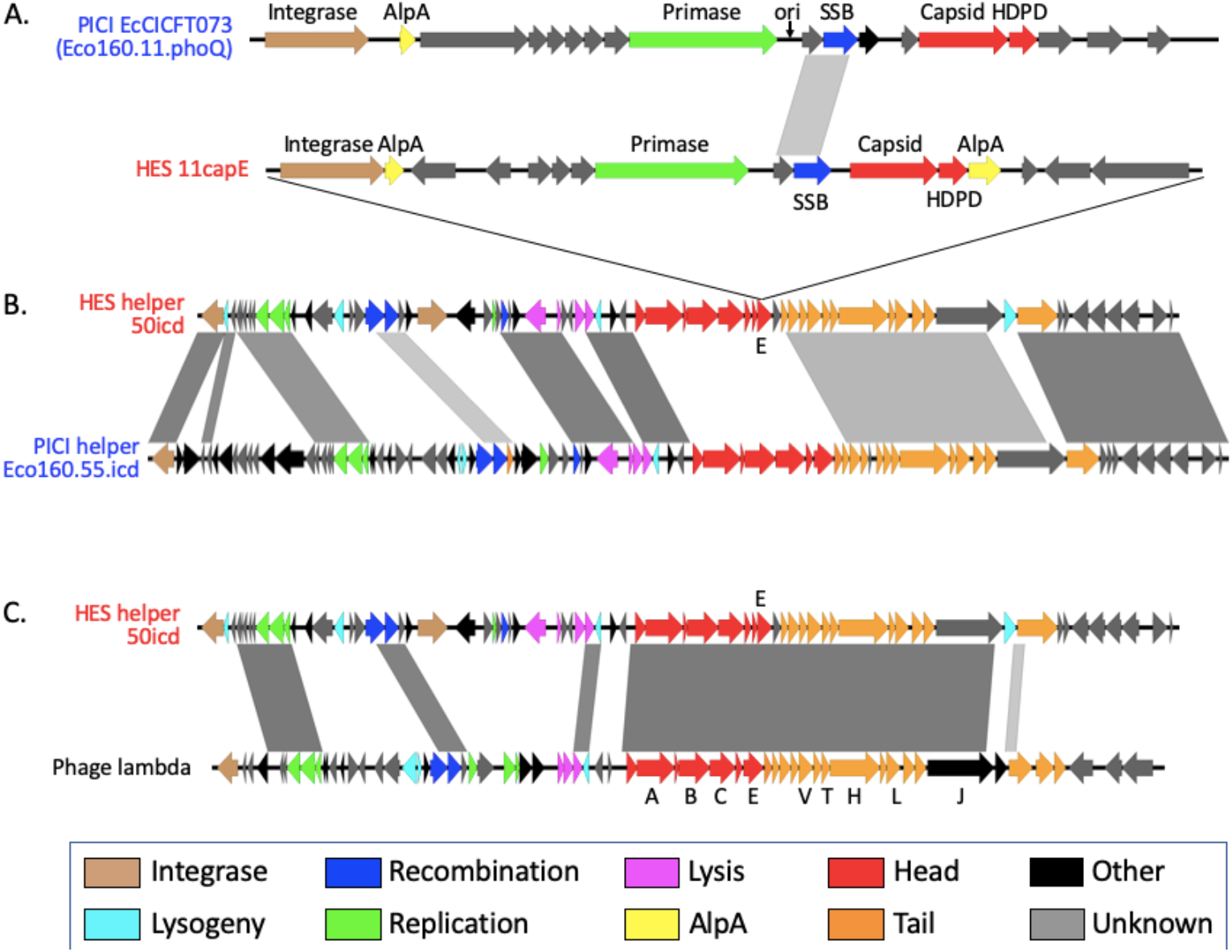
Similarity of the two components of 61icd to a PICI/helper pair. **A**) The 11capE component of 61icd shows similarities with model PICI EcCICFT073 in size, gene content and order, and in some nucleotide sequence regions (parallelograms). The 50icd prophage component shows extensive similarity to the PICI helper prophage (**B**) and to phage lambda (**C**).

### Induction: 61icd excision, replication, and 11capE excision

Induction is a key stage in the life cycle of a temperate phage, a stage where a satellite and helper may interact most^11^. The HES:helper lysogen Eco567 was treated with a standard prophage inducer, the DNA-damaging agent mitomycin C, and the status of the attachment sites of 61icd and 11capE was monitored over time, using two quantitative methods, qPCR and read counting from deep sequencing^22^ (Fig. 3, A and B). Of special interest are the products of excision, *attB* (the chromosomal scar) and *attP* (the circularization junction of the mobile element), while the *attL* and *attR* pairs of the unexcised lysogen are used in normalization. Reads from deep sequencing confirm precise excision of both 11capE and 50icd, producing their TIGER-predicted *attB* and *attP* sites, and likewise for another predicted Eco567 prophage, 41torS.

**Figure 3.**
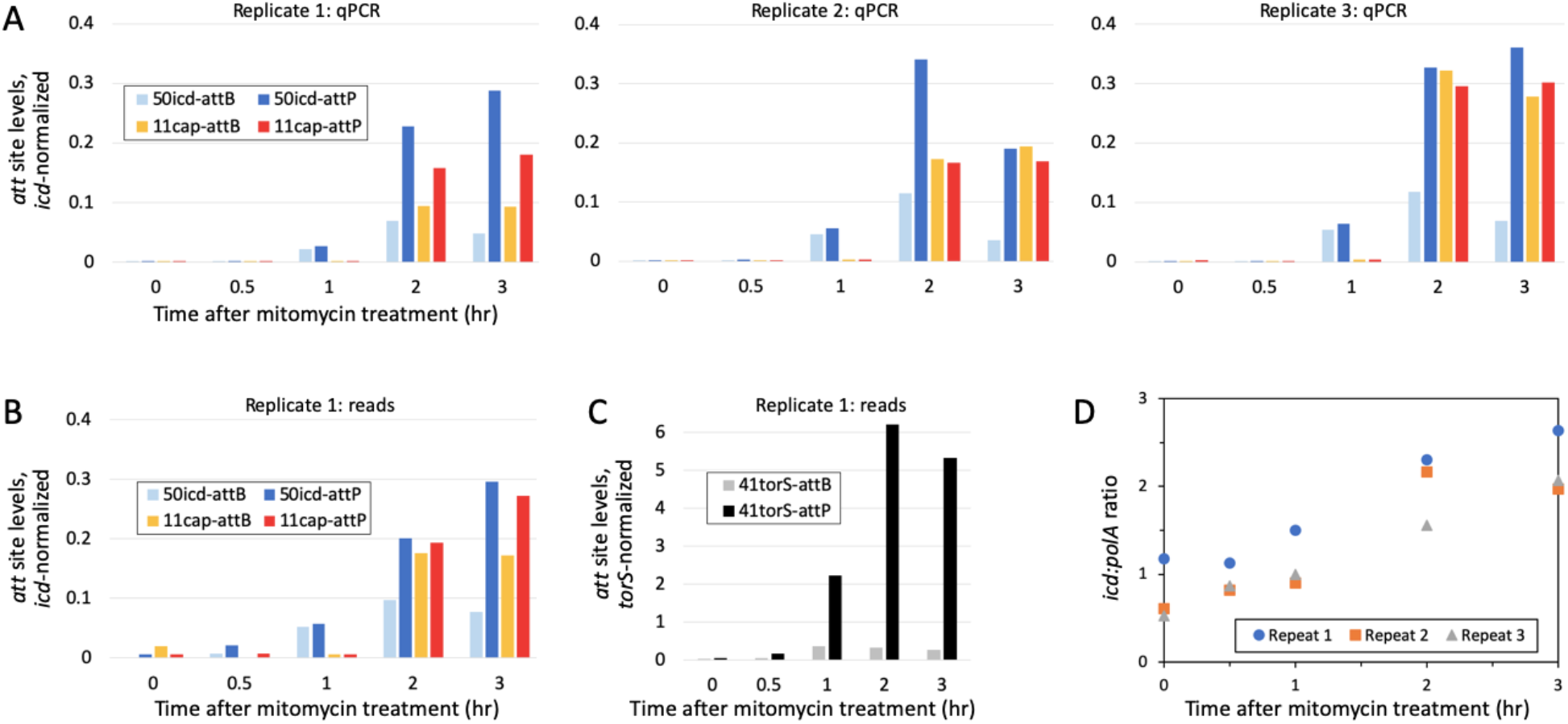
Monitoring recombinant attachment site (*attB* and *attP*) levels after induction. **A**) Three independent replicates of a mitomycin prophage induction experiment were monitored by qPCR. Counts of *att* sites are normalized to those of the *icd* gene as described in Materials and Methods. B) Replicate 1 was subjected to deep sequencing and read counts were taken for each *att* site, normalizing as in A. **C**) Deep sequencing *att* count data for 41torS, normalized to the *torS* gene. Note the 15-fold y-axis increase relative to B. **D**) In situ amplification of *icd* upon induction, normalized to the distant housekeeping gene *polA*, measured by qPCR.

The *icd attB*, representing derepression and excision of 61icd, is detected at 1 hr post-treatment, at levels corresponding to induction in 2-5% of the *icd* genes, rising to a maximum of only 7-11% at 2 hr, and dropping back to 3-7% at 3hr. We interpret this late drop of normalized *attB* as overgrowth by the much larger number of healthier uninduced cells. The separate prophage 41torS is more strongly induced, in up to 35% of *torS* genes (Fig. 3C). No reads corresponding to *attB* or *attP*, indicating excision, were found for a third TIGER-predicted prophage, 45proP.

Levels of *icd attB* and *attP* would be nearly equal if excision were the only event to occur. Instead we observed an excess of *attP* relative to *attB*, evidence that the excised helper/composite replicated faster than the chromosome. Ratios of *icd attP:attB* were only 1.1-1.2 by 1 hr, but rose to 2.1-3.3 by 2 hr and to 3.8-6.0 by 3 hr, indicating post-excision replication of the helper/composite. By contrast, excised 41torS replicated even faster, reaching an *attP:attB* ratio of 20.

11capE excision was not detected until the later timepoint of 2 hr, when substantial replication of the 61icd composite had occurred. The ratio of the 11capE *attB* to the *icd attP*, a measure of the extent of excision of 11capE, was 42-98% at 2 and 3 hr. Experimental repeat 2 showed a delay in excision relative to replication, with an excision yield of ~50% at 2 hr and ~100% at 3 hr.

11capE *attP* and *attB* levels were nearly equal in two of the three experimental repeats. This suggests either that neither of the separated 11capE and 50icd components replicated further after satellite excision, or that they both replicated at equal rates. In the other repeat, the *attP:attB* ratio was at 1.6-1.9, suggesting moderate post-excision replication by 11capE.

In addition to replication of 61icd after excision, there was amplification of unexcised 61icd, evidenced in a steady increase (2.2-3.9 fold over 3 hr) in the ratio of the *icd* gene to the distant *polA* gene (Fig. 3D). This comports with in situ pre-excision replication observed for P2 and other temperate phages^24–27^.

We could not detect plaque-forming units in Eco567 induced lysates, either on lawns of *E. coli* MG1655 or of Eco567 from which we had precisely deleted 61icd (data not shown).

### Additional HESs

We searched for more examples of 11capE by querying enterobacterial genome databases with its integrase sequence. As expected we found more instances, totaling 201 of similar elements in the same site in homologs of the 50icd capsid E gene. Surprisingly, the related integrases also led to 290 instances of HESs in other phage late gene sites, totaling 14 sites in nine late genes (Fig. 4). These 14 site types are standardized using protein profiles of the late proteins, named according to their lambda cognate gene, the protein profile encompassing their attachment site, and the location of the central codon in the attachment site within that profile. For example, 11capE is inserted in the 80^th^ codon of the Pfam profile Phage_cap_E, in a cognate of phage lambda gene E, so we term this site E:Phage_cap_E:80. A possible fifteenth HES site (B:Phage_portal_2:21) could only be guessed, based on late gene coding sequence upstream of the *int* gene; its *attR* could not be located. We can describe our effort as a systematic survey, not for all HESs, but for HESs among Enterobacterales with close relatives of the 11capE integrase. It is very likely that there are more HESs in genomic databases; several of those found here were close to our integrase match cutoff score.

**Figure 4.**
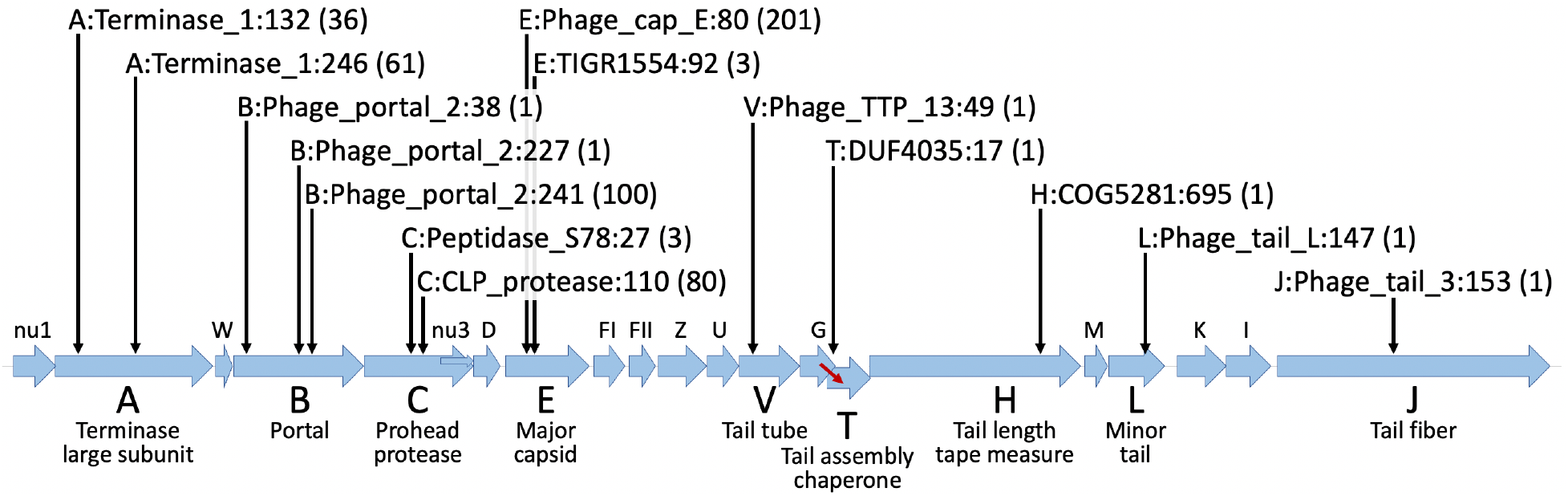
HES integration sites in prophage late genes. A 19-kbp region of phage lambda late genes is shown. Each HES was found in a homolog or functional cognate of one of these late genes, oriented such that the key HES genes (integrase, AlpA, and primase) have the same (rightward) orientation as the late gene. The labels for the 14 detected HES sites have three parts: the lambda gene cognate, the protein family sequence profile that covers the integration site, and the position of the site within that profile; the number of HESs found at the site is in parentheses. The T gene has no start codon; it is only expressed upon a translational frameshift (red arrow) from the overlapping G gene.

For 209 HESs, the surrounding helper prophage was mapped; in one case (Supp. Figure 1D) the same helper prophage (Ana29.79.guaB) contained two HESs at separate late gene sites (A:Terminase_1:245 and V:Phage_TTP_13:49). For 24 additional HESs (all C:CLP_protease: 110), the target late gene was not within a helper prophage, but instead mapped within a PICI itself integrated into a tRNA-Met gene, forming a 20.6-kbp composite. Other C:CLP_protease:110 HESs are in prophages, so the integration into PICIs may be accidental, but could nonetheless reveal interesting biology.

Of satellites mapped to chromosomal sites, 28 were related to a HES group, based on integrase phylogeny (see below) and confirmed by alignment of the *attL* and *attR* sequences (Supp. Files 1 and 2). We interpret these as the result of off-target integration events by HES integrases, much like lambda and other phages can integrate into secondary chromosomal *attB* sites if the normal *attB* is unavailable^28^.

The remaining chromosomal satellites were found at one of two sites, 2642 in the *fis* gene and 49 in the *hpt* gene, representing two new subfamilies of non-HES PICIs. Their integrases matched to 11capE just above the cutoff. A tandem pair of PICIs was noted at the *fis* gene in two genomes (Supp Table 1).

Summarizing the categories of the 3452 mobile genetic elements newly mapped here, we found 491 HESs, 28 off-target HESs, 2701 GN-PICIs (at *fis* or *hpt* loci), 208 helper prophages surrounding HESs and 24 GN-PICIs surrounding HESs.

### Integrase phylogeny

All the new HESs, PICIs and prophages had a primary tyrosine integrase gene near a terminus of the element, and some contained secondary or tertiary internal *int* genes. A phylogenetic tree was prepared for all these integrases, together with the integrases of the 48 unmapped satellites and several reference integrases (Fig. 5). The unifying principle for integrase clades is the use of the same *attB* site. Perhaps unsurprisingly (since HESs were discovered based on integrase similarity), the primary HES integrases form a distinct clade. The HES clade constitutes a branch within a larger clade of mostly PICIs and satellite P4. Some helper integrases also fall in this PICI group, but most helper integrases are from a larger subtending group with integrases from phages lambda and P22. Helper prophages use a variety of chromosomal *attB* sites, and HES types can be mixed within helper clades using the same *attB*. The integrase tree confirms the 34 *attB* guesses for the cases where *attR* could not be identified, and suggests origins for the 9 fully unmapped cases. The integrase of the C:CLP_protease:110 HESs appears the most promiscuous, accounting for 24 of the 28 off-target events. Allowing for these off-targets and for the secondary integrase encoded in nine HESs, each of the 14 site-groups of HESs were coherent (monophyletic) by integrase phylogeny, except for the two slight splittings indicated. The 14 site-groups were further supported by *attL* and *attR* sequence alignments (Supp. File 1). The integrase for the HES guessed to specify a fifteenth site, B:Phage_portal_2:21, formed an isolated branch distant from those of the other three B:Phage_portal_2 sites. In two cases, a secondary HES integrase gene was found to be closely related to that of HES’s helper prophage.

**Figure 5.**
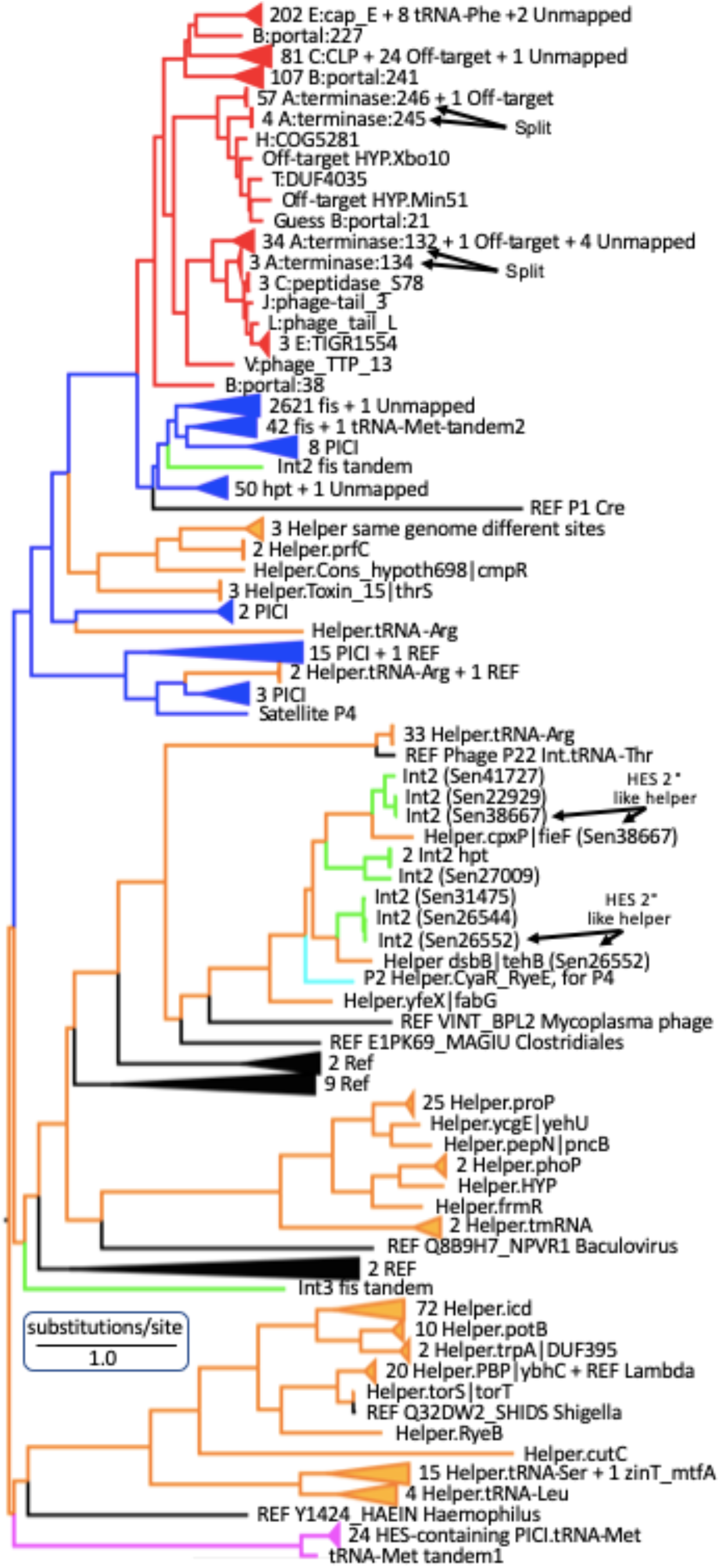
Integrase phylogeny. The primary integrase of each HES (red), reference or newly found GN-PICI (blue), helper prophage (orange) or surrounding PICI (magenta), unmapped elements, the secondary integrases encoded by some of these satellites (Int2, green), and reference integrases (black) were aligned to build a maximum-likelihood tree. Each *attB* that had been guessed for an unmapped element was confirmed by the tree. nine fully unmapped elements are indicated. For each helper clade the *attB* site specificity is indicated.

### Attachment sites

Tyrosine integrases (the only type used by the GIs studied here) tend to enforce sequence identity between *attP* and *attB* at the typically 7-bp segment where strand exchange occurs^29^. This core of the *att* site identity block can extend substantially further (up to ~250 bp), when an integrase specifies an internal site well within the target chromosomal gene, as occurs at *icd* and many tRNA genes. In these cases, the *attP* site carries a replacement sequence such that the target gene is still functional after integration. TIGER identifies such extended identity blocks by comparing *attL* and *attR* from the query genome to the uninterrupted *attB* of a reference genome, but cannot determine the crossover core within it. Comparison of diverse sequence instances can help localize the core, narrowing the crossover site within the *fis* gene *attB*, despite often appearing within long identity blocks, to a sequence that is typically TTCATGCC, and the site within the *hpt* gene to AAACATA.

The integration event in Eco15648.11.E:Phage_cap_E:80 left an extra adenosine residue at both *attL* and *attR*, shifting the frame of the late gene target. Long duplications of the flanking target gene were observed for the *att* sites of two HESs (Eco15150.12.Phage_cap_E_154 and Eco6102.18.E_Phage_cap_E_39), suggesting improper recognition of *attB* during integration. Some drift of the central codon of the attachment site identity block was observed, perhaps due to idiosyncratic mutations near the crossover segment within the identity block; for example one group ranges from codons 240-243 on the Pfam profile Phage_portal_2. Such *attB* drift among very similar integrases was also observed for reference PICIs integrated into the yicC gene, where drift was so extreme that the identity blocks do not overlap.

### Gene content

Proteins of the HESs, PICIs and helpers discovered here, together with reference GIs (3484 total) were clustered into families based on similarity scores. This provided a standardized gene profile to each GI, reducing the GIs to 646 unique genotypes (218 for HESs and their off-targets, 259 for the PICIs, one for P4, and 168 for the prophages). This reduction acts to correct redundancy due to over-representation in the genome databases of the genera *Escherichia, Salmonella* and *Klebsiella*.

We note some peculiar satellites: 1) One group is integrated in tRNA-Phe genes, despite having an integrase extremely similar to that specifying the E:Phage_cap_E site; these lack a Primase and uniquely encode a Resolvase (Supp. Figure 1C). 2) The HES-invaded PICIs shared a single genotype, bearing a CLP_protease gene containing the HES *attB* (Supp. Figure 1B). These PICIs are integrated into tRNA-Met genes, with integrases that are phylogenetically quite distinctive (bottom of Fig. 4), and also lack an *alpA* gene. 3) For Eco6102.18.E_Phage_cap_E_39, a 695-bp segment containing the HDPD gene is repeated at the left portions of both *attL* and *attR*, and the first 102 nt of that segment is repeated before (and overlapping by 4 bp) the *attR* copy. 4) A tandem satellite pair was also found within a tRNA-Met gene, with one integrase similar to that of the HES-invaded PICIs (Supp. Figure 1G). Normally the three *att* sites defining a tandem pair have similarities, but in this case, the terminal *att* sites are adapted for tRNA-Met and not related to the internal *att* site which is a *fis* PICI *attP*. Thus it is difficult to envision how the two units would join by integrase activity. Perhaps a circular *fis* PICI entered the tRNA-Met PICI by homologous recombination within the long segment that is duplicated in the tandem.

We observed four ordered zones among HESs and PICIs in the upstream to downstream direction: Integration, Bro, Replication, and Late. In the Integration category, with Integrase proteins, we include HTH_17 and the multifunctional Phage_AlpA, known as excisionases controlling the directionality of integration^18^. The Bro zone is dominated by domain-swapping proteins of the Bro network^30^. Their structural similarity to homing endonucleases has been noted, suggesting possibilities for DNA recombination activity. A member encoded by *Vibrio* satellite PLE has been shown to act as an immunity nuclease^31^. The dynamism of these Bro network genes among the satellites required a specialized annotation approach (Methods) that provides further evidence of domain swapping. Also encoded within some Bro zones are immunity RNAs; In P4, the RNA repressor CI is encoded within a Bro network gene and controls early transcriptional termination by binding to a nearby RNA site, seqA^32^. Two Rfam profiles (C4 and isrK) can detect CI and seqA relatives^33^, and these hit several of our GN-PICIs and HESs. The Primase, SSB and DUF5906 proteins, whose genes colocalize along with mapped origin of replication sequences^6^, were placed in the Replication category.

The downstream Late zone, like the Bro zone, is another dynamic region of HES/PICIs, containing diverse prophage late genes, as identified by co-oriented gene clusters from our own prophage set (Supp. Figure 2). Our late gene categorization system was conservative and did not find late genes among the most divergent satellites, the reference PICIs of the Pasteurellaceae. This may reflect bias toward Enterobacteriaceae late genes by our system; indeed capsid genes have been identified in most of these Pasteurellaceae PICIs^6^, in our Late zones. Long late gene clusters found in some satellites perfectly match those in our prophages, suggesting direct capture into satellites of late gene clusters from their helpers (Fig. 6).

**Figure 6.**
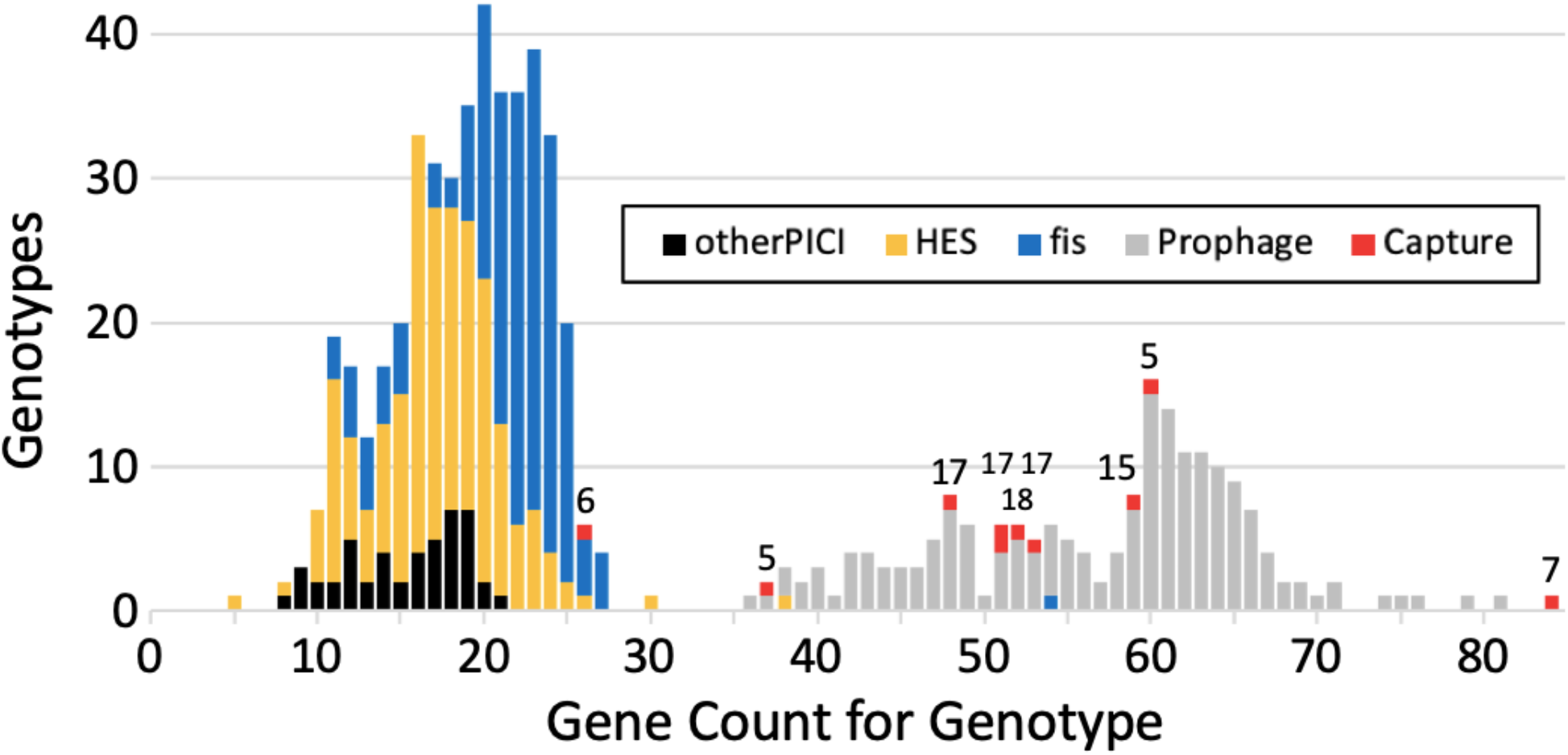
Capture of prophage gene clusters in longer HESs and PICIs. The count of protein genes was taken for the 646 GI genotypes. The longest gene cluster shared with any of the prophages was taken for each PICI and HES genotype; 10 of these (Capture) were prophage gene clusters of 5 or more genes (numbers denote the cluster sizes).

Bro network protein genes, such as P4 *eta*, can be the scene of multiple regulatory activites in satellites^34^. The transcript from a constitutive promoter internal to *eta* is quickly terminated due to RNA interactions at the CI and seqA sites; processing of this transcript produces the stable immunity RNA CI. Homologs of CI and seqA were found in several of the HES/PICI Bro genes, suggesting that their Bro zones provide immunity functions. A truncated form of Eta (Kil, which functions in cell killing) can be translated upon occasional terminator read-through. Full-length Eta cannot be translated until an upstream late promoter is activated; *eta* is translationally coupled to the upstream *alpA* homolog. Eta translation prevents termination, so that the late promoter reads through the full *eta* gene and downstream genes.

Bro network proteins often have gene regulatory functions; they can mix domains related to nucleases with other domains that have a distinct DNA-binding fold^30^. One example, CapR of the *Vibrio* satellite PLE represses the capsid gene of its helper phage by binding its promoter −35 region^31^. CapR fits into the Bro network by its similarity to domains in proteins with a Bro network nuclease domain. A protein (MJY48377.1) of the HES Eco11232.16.A_Terminase_1_246 contains the HNH_3 nuclease domain, linked to the AP2 domain and the Bro segments termed here b60, b155, b3, and b22.

## DISCUSSION

We describe a new satellite/helper relationship where the satellite genome lies embedded within its helper prophage genome. Analysis of induction indicates a phase, after excision of the 61icd HES/helper composite from its integration site in *icd*, when the entire composite replicates. To the extent that the helper 50icd component drives this replication, the HES 11capE is passively replicated as a segment within the larger composite, an example of cis-regulation not possible in standard satellite/helper pairs. However 11capE has its own primase and origin of replication, so may well also contribute to replication of itself and the composite.

Additional interactions between the two components of 61icd can be proposed. Certainly the 50icd capsid gene cannot be expressed until the gene is restored by excision of 11capE. This likely reduces 50icd phage yields, in line with other model satellites (reviewed in ^11^) known to partially inhibit their helpers. We have discussed how 50icd serves as a helper for 11capE DNA replication, and suspect that 50icd is also the helper for three other functions: 1) detecting the SOS response signals for induction through its cI repressor, 2) providing the structural proteins and packaging machinery to make 11capE virions, in line with other lambdoid prophage helpers of PICIs, and 3) activating 11capE gene expression in the helper late phase. All of the non-hypothetical 11capE genes are unidirectional, in the same direction as the 50icd capsid gene and its surrounding late genes. It can be hypothesized that late transcription of 50icd will continue into integrated 11capE, transcribing its integrase, capsid and other genes, and will do so at a time when much replication has already occurred. It should be noted that another prophage in these cells, 41torS, is induced earlier, in a larger fraction of the cells, and with faster replication, which may modulate the 61icd activities observed here.

Further analysis was hampered by an inability to detect the expected virions of 50icd, even on lawns of its source strain cured of 61icd, though there may be future paths forward. An interesting remaining question is whether the entire 61icd composite may be packaged into some virions, delivering helper and satellite together into a new host. Otherwise, co-delivery to a new host can occur the usual two-virion way, with low levels of helper phage virions released together with satellite virions upon lysis.

Surprisingly, a search based on integrase similarity revealed many additional HESs integrated into diverse prophage late gene sites. This constitutes evidence for a tropism of an integrase clade toward integration sites in prophage late genes. It will be of interest to learn whether mechanistic peculiarities of this integrase subclade account for this tropism. It is possible that the neighboring *alpA* gene, encoding a tightly conserved excisionase/transcription factor, contributes to the tropism. Although we found HESs at 14 *att* sites in nine late genes, 97% of them were at five sites in the four genes A, B, C and E in a core region occupying only 6.5 kbp of phage lambda. The most frequent site was in the capsid gene E. This result may be biased, because our search was launched with one of those satellites, yet it has additional significance; capsid genes are the most common late phage gene in GN-PICIs. We find that HESs tend to have captured more phage late genes than the GN-PICIs, which may be facilitated by their localization and recombination activity in these same prophage regions. Phage late genes more likely to be captured in the HES are those closest to the HES integration site. This provides a backup intact copy that may functionally replace the interrupted prophage gene even before HES excision. This redundancy is not essential; knockout of a capsid gene in a PICI was tolerated^6^. The prophage/satellite exchange observed here contrasts with the situation in *Streptococcus*, where prophage/satellite pairs are widespread, but have little genetic exchange^35^. The HES champion is *Arsenophonus nasoniae* FIN, whose genome has four helper prophages each bearing a slightly different B:Phage_portal_2:241 HES, with one of these helpers additionally bearing a second HES at a different late gene (Supp. Figure 1D).

When an integrase targets a site within a tRNA or protein gene, the *attP* usually carries within itself a replacement sequence from the target gene, positioned such that the postintegration recombinant target gene is still functional. There are exceptions to this rule, that can usually be understood as cases of regulated gene integrity, where disruption of the target gene upon island integration serves to regulate cellular processes such as sporulation and DNA replication fidelity^16, 36^. HESs disrupt prophage late genes, but do not carry a replacement sequence in *attP*. Thus HESs appear to cause regulated gene integrity, in which the regulation serves the HES, rather than the cell or the prophage.

HESs serve to remind that integrative elements can target not only chromosomal genes but other mobile elements. Interestingly, PICI EcCIDi14 is in a IS110 transposase gene^6^. By gene content, the HESs fall squarely among the PICIs, although the term PICI is less apt for them; more than “phage-induced”, HESs are phage-embedded, and are thus not “chromosomal islands” but islands of their helper prophages.

11capE is presumed to be packaged by the *cos* machinery of the helper, as has been presumed for EcCICFT073^7^, which likewise lacks *terS;* a 258-bp sequence is shared with 90% identity between 11capE and 50icd, at the same site in 11capE where *cos* sites have been identified in other PICIs, and at the location in 50icd that maps to the lambda *cos* site.

Discovery of GIs, including prophages and PICIs, is a necessary capability to fully characterize the bacterial genomic landscape. Computational tools for detecting integrated satellites have been lacking^37^, but this work shows that TIGER/Islander software is suitable for fully automated and precise mapping, in spite of their small sizes and ability to use rare integration sites. Scale-up of our software to all available bacterial genomes is expected to identify numerous satellites in other bacterial groups.

PICIs and HESs exemplify different integration strategies. PICIs typically target conserved housekeeping genes, where they may wait for arrival of the helper if it is not already there. In contrast for HESs, when the target site is available (helper is present) it favors an especially profitable HES location that can provide several cis-regulatory advantages. However, the target site will be more often absent for a HES than for a PICI, because the HES helper may be absent or its target late gene may be more subject to change than a housekeeping gene. This may explain our finding of several off-target HES integration events.

Inspection of annotations showed that many of the HES helpers appeared truncated in their chromosomal setting; such potentially abandoned HESs may retain functions allowing activation when a new helper enters the cell. The local source of phage genes in the helper may facilitate the recombinational shuffling of phage genes into HESs.

HESs are not particularly widespread; we estimate that among the *E. coli* genomes, where chances were best for finding 11capE-related HESs, only ~5.4% of genomes contain a HES (this calculation accounted for the high fraction, 56%, of 11capE *int* hits that fell in a region unsuitable for HES detection due to contig truncation or ambiguous bases).

The HESs, PICIs, P4, and the *Vibrio* PLEs are rich in Bro network proteins and will provide an excellent system to better characterize their interesting domain-swapping evolutionary history.

## ACKNOWLEDGEMENTS

This work was supported by the Laboratory Directed Research and Development program at Sandia National Laboratories, which is a multimission laboratory managed and operated by National Technology and Engineering Solutions of Sandia LLC, a wholly owned subsidiary of Honeywell International Inc. for the U.S. Department of Energy’s National Nuclear Security Administration under contract DE-NA0003525.

We thank Alfredo G. Torres for the kind gift of *Escherichia coli* NRG857c.

## AUTHOR CONTRIBUTIONS

D.T. and C.M.M. performed experiments, data analysis, results interpretation and writing. K.P.W. did data analysis, results interpretation, writing and acquired funding. All authors revised and approved the manuscript.

