## Supplemental File 1 for "An integrase clade that repeatedly targets prophage late genes, yielding helper-embedded satellites"

**SUPPLEMENTARY MATERIALS**

Supplementary Files 1 and 2. Attachment sites. The *attL* (File 1) and *attR* (File 2) sequences of each model GI (except the tandem Eae41.tRNA-Met satellites) are shown, not fully aligned but simply shifted to optimize alignment of the identity block (lower case) within in each group. The 14 HES groups with their off-site relatives are at the top, followed by PICI groups then prophage groups. All tRNA sites used are at the bottom, with the complete tRNA sequence in *attL* aligned based on secondary structure, and the *attR*s aligned to emphasize damage at the 3′ portions of the Pro, Thr, and Ile2 tRNAs [Williams KP. Integration sites for genetic elements in prokaryotic tRNA and tmRNA genes: sublocation preference of integrase subfamilies. Nucleic Acids Res 2002;30:866-875]. Sites for the tandem Eae41.tRNA-Met satellites are omitted here.

Supplementary Table 1. Study genomic islands (separate file)

Supplementary Table 2. Primers used in this study

| **Oligonucleotide** | **Sequence** | **Use** |
|---|---|---|
| 143.11capE.CJ.L | CGGCCACCTGCTAACCTGTA | Identifying 11capE attL, attP |
| 144.11capE.CJ.R | AGTTTTGATGCTCGTTTGATATCCC | Identifying 11capE attR, attP |
| 145.50icd.CJ.L | GCGATCTCTGTCAGAACGGT | Identifying 50icd attL, attP; qPCR |
| 146.50icd.CJ.R | GAGACACATAAGGCCTCGCA | Identifying 50icd attR, attP |
| 147.Eco567.icd.DJ.L | ACCCGAATACCGGCAAAGAG | Identifying 50icd attL, attB |
| 148.Eco567.icd.DJ.R | AAGAGGCCGCATTGCTTCAT | Identifying 50icd attR, attB |
| 149.Eco567.capE.DJ.L | GCCGGAACGGCAATCAGC | Identifying 11capE attL, attB |
| 150.Eco567.capE.DJ.R | CAGCGCGTAGGCTTCGATATCG | Identifying 11capE attR, attB |
| 319.11capE_Q_CJ_L | AGGAAAATTCAGTGAAAGCGCC | Quantifying 11capE attL, attP |
| 320.11capE_Q_CJ_R | TGGTTGCATCGTGTCATCCT | Quantifying 11capE attR, attP |
| 321.11capE_Q_DJ_L | AACATGGCGCTGTACGTTTC | Quantifying 11capE attL, attB |
| 322.11capE_Q_DJ_R | GCCTGCATCTCTTCGACCTG | Quantifying 11capE attR, attB |
| 355.Eco567.icd_DJ_L_qPCR | ATGCCGGACAGGACAAAGTA | Quantifying 50icd attL, attB |
| 356.50icd_CJ_R_qPCR | CGAACGTCAATGAAATCAAACGGT | Quantifying 50icd attR, attP |
| 357.Eco567.icd_DJ_R_qPCR | GCTCCCCATAAATAATCACCAGAC | Quantifying 50icd attR, attB |
| 366.Eco567_Q_polymerase_L | TAACCTGGTCGGGCTTTCTT | Quantifying DNA polymerase I |
| 367.Eco567_Q_polymerase_R | GCTTCTTCGAGGGCAATCTG | Quantifying DNA polymerase I |

Supplementary Table 3. Probes used for AttCt to detect predicted mobile elements and housekeeping gene *polA* (DNA polymerase I).

| **Element Name** | **attB** | **attP** | **attL** | **attR** |
|---|---|---|---|---|
| *polA* | N/A | N/A | TAACCTGGTCGGGC | GCTTCTTCGAGGGC |
| 61icd | GATGCTGCGCCACA | ACTGCTGCGCCATA | GATGCTGCGCCATA | ACTGCTGCGCCACA |
| 11capE | AAACCCAAGCATG | CATGTAACCCAAGC ACTTAG | AAACCCAAGCACTT | TAACCCAAGCATGA |
| 41torS | GCACTTTAGGTGAA AAAGGTTGAGT | ATCCTTTAGGTGAA TAAGTTGTATA | GCACTTTAGGTGAA TAAGTTGTATA | ATCCTTTAGGTGAA AAAGGTTGAGTC |
| 2emrE | CACCACATTAAAAA TAATTTATTTTTAA ACGACTAAAATAGG TT | GCCCCCATTAAAAA TAATTTATTTTTAA ACGACTAAAATATG GA | CACCACATTAAAAA TAATTTATTTTTAA ACGACTAAAATATG GA | GCCCCCATTAAAAA TAATTTCTTTTGAAT CGAGTGAAATAGGT T |
| 48F | CAGGGGATTGAAA ATCCCCGTGTCCTT GGTTCGATTCCGAG TCCGGGCACCAAAT | TCAGGGATTGAAAA TCCCCGTGTCCTTG GTTCGATTCCGAGT CCGGGCACCACTA | CAGGGGATTGAAA ATCCCCGTGTCCTT GGTTCGATTCCGAG TCCGGGCACCACTA | TCAGGGATTGAAAA TCCCCGTGTCCTTG GTTCGATTCCGAGT CCGGGCACCAAAT |
| 7betA | AATACTGGATATGA AGCATGA | GATTGTCACGATTT ATAGCCT | AATACTGGATATTT ATAGCCT | GATTGTCACGATGA AGCATGA |
| 45proP | GCTTGAATTATGGA CTTCCAGTTAT | CTGTGAATTTATGG ATTTCAAGAGT | GCTTGAATTTATGG ATTTCAAGAGT | CTGTGAATTATGGA CTTCCAGTTAT |
| 4fimD | ATGCAGGGCGTTTA CGCGACA | GTGCGTGGCGTTTT TACGGCA | ATGCAGGGCGTTTT TACGGCA | GTGCGTGGCGTTTA CGCGACA |

Supplementary Figure 1. Variant satellite configurations. A) Standard HES in a helper prophage. B) HES in a PICI. C) PICI in a tRNA gene, though using the same integrase subclade as the HES in panel A. D) Helper with two HESs. E and F) Simple tandems in *fis*. G) Complex tandem between a tRNA-Met PICI and a *fis* PICI, where the two share a long duplicated region.

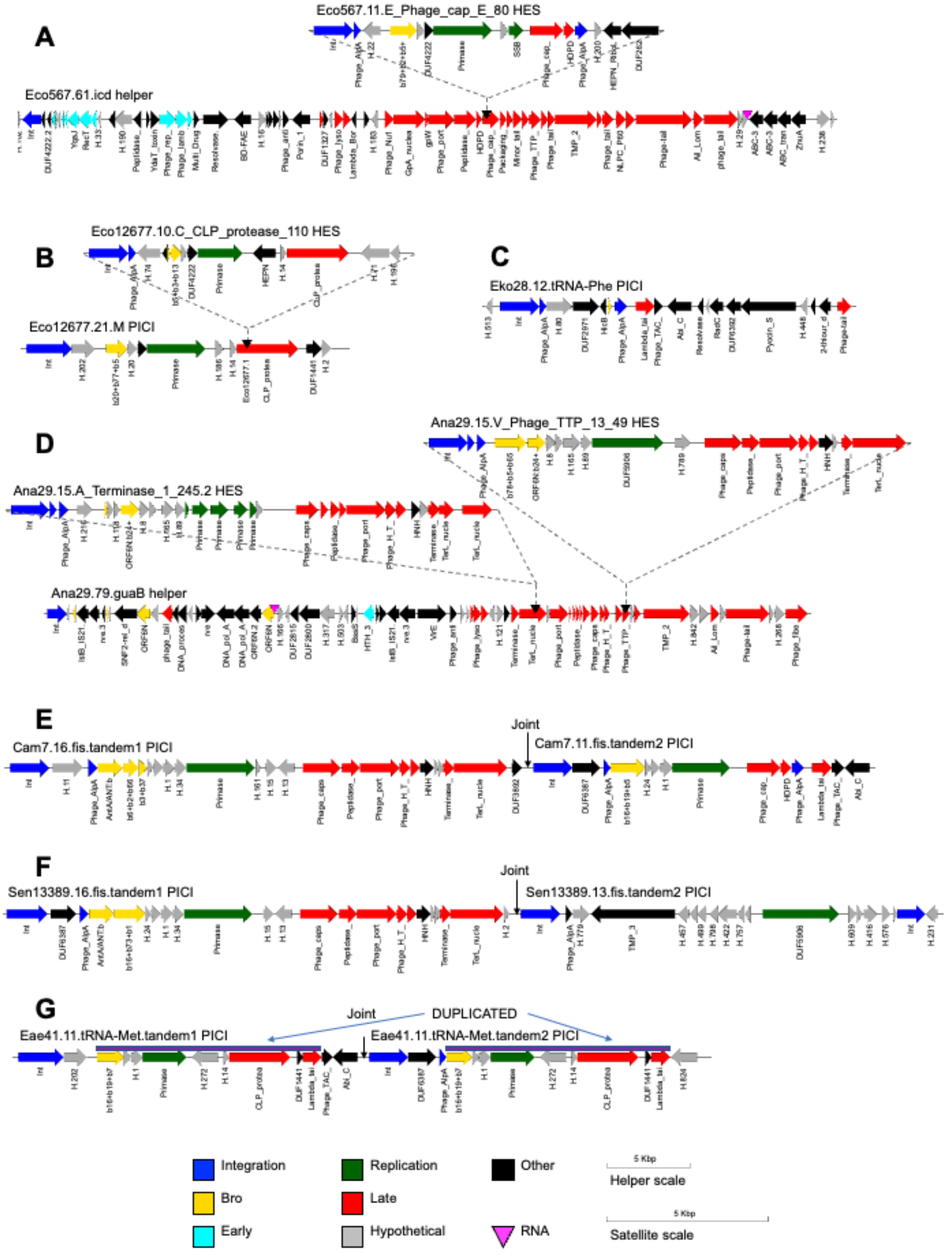

Supplementary Figure 2. Late/early genes in co-oriented strings. The 168 gene profiles for prophages were split into continuous strings of co-oriented genes, collecting those containing a holin/tail seed family as "late strings". Based on transcription profiles for reference phages lambda and P2, non-seed families were labeled "Known Early" or "Known Late" and the remainder "Test". For each family the ratio, presence in lates strings : presence in profiles, was computed.

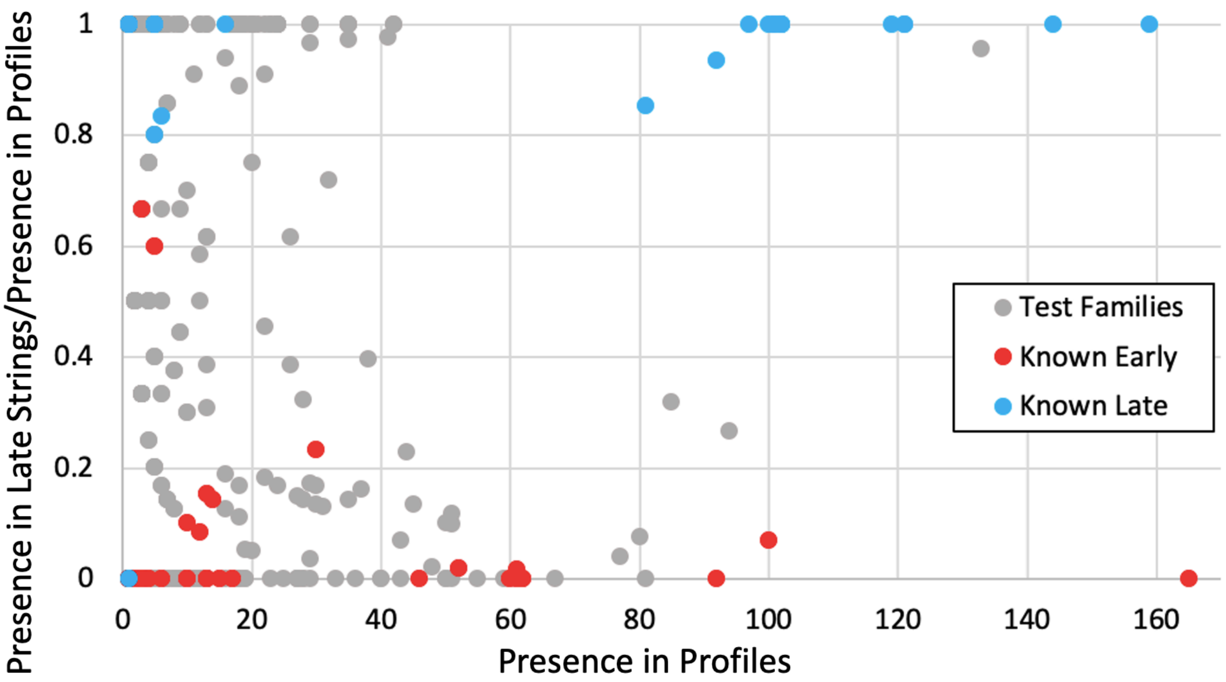


Supplementary Figure 3. PCR products used as standards in qPCR experiment. Agarose gel showing PCR products for the eight attachment sites of interest and the *polA* housekeeping gene (HK). These products were used as standard templates after being diluted to $1x10^{-2}$ ng, $1x10^{-4}$ ng, $1x10^{-6}$ ng, and $1x10^{-8}$ ng. They were amplified and standard curves were generated by plotting Ct value with respect to log starting concentration in nanograms.

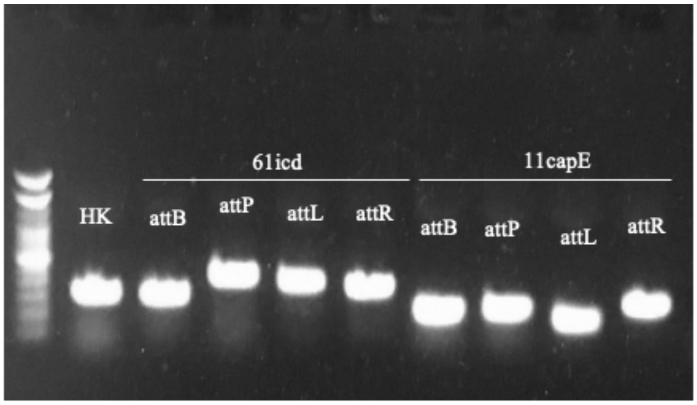
